# Model design for non-parametric phylodynamic inference and applications to pathogen surveillance

**DOI:** 10.1101/2021.01.18.427056

**Authors:** Xavier Didelot, Lily Geidelberg, The COVID-19 Genomics UK (COG-UK) consortium, Erik M Volz

## Abstract

Inference of effective population size from genomic data can provide unique information about demographic history, and when applied to pathogen genetic data can also provide insights into epidemiological dynamics. The combination of non-parametric models for population dynamics with molecular clock models which relate genetic data to time has enabled phylodynamic inference based on large sets of time-stamped genetic sequence data. The methodology for non-parametric inference of effective population size is well-developed in the Bayesian setting, but here we develop a frequentist approach based on non-parametric latent process models of population size dynamics. We appeal to statistical principles based on out-of-sample prediction accuracy in order to optimize parameters that control shape and smoothness of the population size over time. We demonstrate the flexibility and speed of this approach in a series of simulation experiments, and apply the methodology to reconstruct the previously described waves in the seventh pandemic of cholera. We also estimate the impact of non-pharmaceutical interventions for COVID-19 in England using thousands of SARS-CoV-2 sequences. By incorporating a measure of the strength of these interventions over time within the phylodynamic model, we estimate the impact of the first national lockdown in the UK on the epidemic reproduction number.

## INTRODUCTION

Past fluctuation in the size of a population are reflected in the genealogy of a sample of individuals from that population. For example, under the coalescent model, two distinct lines of ancestry coalesce (i.e. find a common ancestor) at a rate that is inversely proportional to the effective population size at any given time (Kingman 1982; Griffiths and Tavare 1994; Donnelly and Tavare 1995). More coalescent events are therefore likely when the population size is small compared to when the population size is large. This causal effect of population size on genealogies can be reversed in an inferential framework to recover past population size dynamics from a given pathogen genealogy. This approach to inference of past demographic changes was first proposed 20 years ago (Pybus et al. 2000, 2001; Strimmer and Pybus 2001) and has been fruitfully applied to many disease systems (Pybus and Rambaut 2009; Ho and Shapiro 2011; Baele et al. 2016).

Population size analysis is often performed within the Bayesian BEAST framework (Suchard et al. 2018; Bouckaert et al. 2019) which jointly infers a phylogeny and demographic history from genetic data. Here we focus on an alternative approach in which the dated phylogeny is inferred first, for example using treedater (Volz and Frost 2017), TreeTime (Sagulenko et al. 2018) or BactDating (Didelot et al. 2018), and demography is investigated on the basis of the phylogeny. Although potentially less sensitive, this approach has the advantage of scalability to very large sequence datasets. This post-processing approach also allows more focus on models and assumptions involved in the demographic inference itself as previously noted in studies following the same strategy (Lan et al. 2015; Karcher et al. 2017; Volz and Didelot 2018; Volz et al. 2020). However, some of the methodology and results we describe here should be applicable in a joint inferential setting as well.

The reconstruction of past population size dynamics is usually based on a non-parametric model, since the choice of any parametric function for the past population size would cause restrictions and be hard to justify in many real-life applications (Drummond et al. 2005; Ho and Shapiro 2011). However, even if a non-parametric approach offers a lot more flexibility than a parametric one, it does not fully circumvent the question of how to design the demographic model to use as the basis of inference. For example, the *skygrid* model considers that the logarithm of the effective population size is piecewise constant, with values following a Gaussian Markov chain, in which each value is normally distributed around neighbouring values and standard deviation determined by a precision hyperparameter (Gill et al. 2013). This model can be justified as the discretisation of a continuous *skyride* model in which the logarithm of the population size is ruled by a Brownian motion (Minin et al. 2008). Alternatively, the *skygrowth* model is a similar Gaussian Markov chain on the growth rate of the population size (Volz and Didelot 2018). Both models can be conveniently extended to explore the association between population size dynamics and covariate data (Gill et al. 2016; Volz and Didelot 2018).

The *skygrid, skygrowth* or other similar models can be assumed when performing the inference of the demographic function, and the effect of this model choice has not been formally investigated. Furthermore, these non-parametric models require several model design choices which are often given little consideration in practice. This includes the number of pieces in the piecewise constant demographic function, the location of boundaries between pieces, and the prior expectation for the difference from one piece to another. All of these model design choices may have significant effect on the inference results. Here we propose several statistical procedures to optimise these variables. In particular, the parameter controlling the smoothness of the population size function is usually assumed to have an arbitrary non-informative prior distribution in a Bayesian inferential setting (Minin et al. 2008; Gill et al. 2013), whereas we show here that it can be selected using a frequentist statistical approach based on out-of-sample prediction accuracy. We tested the effect of these procedures on simulated datasets, where the correct demographic function is known and can be used to assess the relative accuracy of inference under various conditions. We applied our methodology to a previously published dataset of *Vibrio cholerae*, the causative agent of cholera. We also analysed a state-of-the-art real dataset and show how our methodology can be used to estimate the impact of non-pharmaceutical interventions for SARS-CoV-2 in England.

## MATERIALS AND METHODS

### Demographic Models

Let the demographic function *N*_e_(*t*) denote the effective population size of a pathogen at time *t*. Let us consider that *N*_e_(*t*) is piecewise linear with *R* pieces of equal lengths *h* over the timescale of interest. Let *γ*_*i*_ denote the logarithm of the effective population size in the *i*-th piece. In the *skygrid* model (Gill et al. 2013), the values of *γ*_*i*_ follow a Gaussian Markov chain, with the conditional distribution of *γ*_*i*+1_ given *γ*_*i*_ equal to:

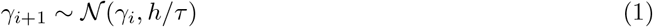

By contrast, the *skygrowth* model (Volz and Didelot 2018) is defined using the effective population size growth rates *ρ*_*i*_ which are assumed constant in each interval and are equal to:

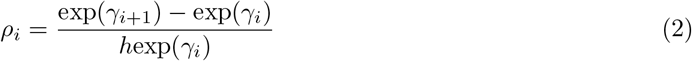

These growth rate values form a Gaussian Markov chain, with:

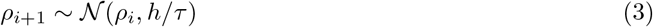

We also define a new model which we call *skysigma* based on the values *σ*_*i*_ of the second order differences of the logarithm of the effective population size:

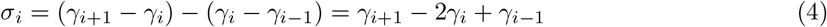

Once again we consider a Gaussian Markov chain in which:

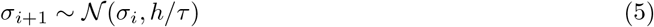

Dependency on known covariate time series can be easily incorporated into these models as previously described (Gill et al. 2016; Volz and Didelot 2018). Let there be a *m* × *p* matrix *X*_1:*m*,1:*p*_ of *p* covariate measurements for each of *m* time points. Ideally these time points would correspond to the *R* + 1 boundaries between pieces of the demographic function, but otherwise linear interpolation can be used to make it so. We model the effect of this covariate data as a modification of the expected change in the demographic variables defined above (*γ*_*i*_, *ρ*_*i*_ or *σ*_*i*_). For example, in the *skysigma* model (Equation 5), the kernel of the Markov chain becomes:

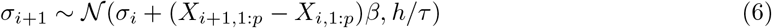

where *β*_1:*p*_ is a vector of coefficients for a linear model of the covariate data on the expected value of the increments. Note in particular that if a term in the *β* vector is equal to zero, then this covariate measurement has no effect on the demographic function, so that to test the significance of covariate requires to test whether the corresponding value in the *β* vector is non-zero.

### Coalescent framework

Each of the models above defines a demographic function *N*_e_(*t*) from which the likelihood of the genealogy 𝒢 can be calculated as briefly described below. Let *n* denote the number of tips in 𝒢, let *s*_1:*n*_ denote the dates of the leaves and *c*_1:(*n*−1)_ denote the dates of the internal nodes. Let *A*(*t*) denote the number of extant lineages at time *t* in 𝒢 which is easily computed as the number of leaves dated after *t* minus the number of internal nodes dated after *t*:

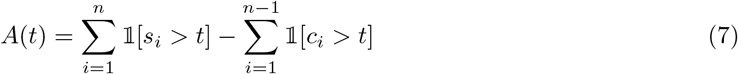

This quantity is important because in the coalescent model, each pair of lineages finds a common ancestor at rate 1*/N*_e_(*t*), so that the total coalescent rate at time *t* is equal to:

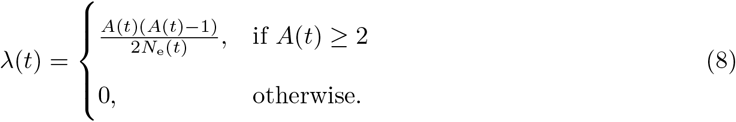

The full likelihood of the coalescent process is therefore computed as (Griffiths and Tavare 1994; Donnelly and Tavare 1995):

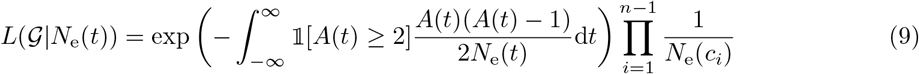

This computation is straightforward for the models considered here where the demographic function *N*_e_(*t*) is piecewise constant.

### Selection of the precision parameter

The demographic models described above (*skygrid, skygrowth* and *skysigma*) all rely on a precision parameter *τ* (also known as the ‘smoothing’ parameter). The value of *τ* controls how much consecutive values of the effective population size will vary when the data is uninformative. The selection of this parameter is therefore shaped by competing aims of optimising the fit to observed data and maximizing explanatory power and avoidance of overfitting. In frequentist statistics, a standard approach to selecting smoothing parameters is to minimize the out-of-sample prediction error. Here, we pursue a *k*-fold cross-validation strategy where genealogical data is partitioned into *k* sets, *k* − 1 of which are used for fitting, and the last one is used for prediction. This procedure is equivalent to maximizing the following objective function:

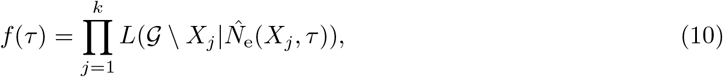

where 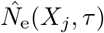 is the maximum likelihood estimates of *N*_e_ on the partial data *X*_*j*_ ⊂ 𝒢 and assuming the precision parameter is *τ* . In this case *X*_*j*=1:*k*_ represents a subset of the sample times and internal node times of the genealogy 𝒢.

This is a standard formulation of the cross-validation method, but the implementation depends on how genealogical data is partitioned. We use the strategy of discretizing the coalescent likelihood (Equation 9) into intervals bordered by the time of nodes (tips *s*_*i*_ or internal nodes *c*_*i*_ of the tree) and/or the *R* − 1 times when the piecewise-constant *N*_e_ changes value. Given *R* − 1 change points, *n* tips, and *n* − 1 internal nodes of G, there are *R* + 2*n* − 3 intervals (*ι*_1_, … , *ι*_*R*+2*n*−3_). Each cross-validation training set is formed by taking a staggered sequence of intervals and collecting the genealogical data contained in each, so that *X*_*k*_ = {*ι*_*j*=1:*R*+2*n*−3_|modulo(*j, k*) ≠ 0}.

### Selection of the grid resolution

Before any of the non-parametric models described above can be fitted, the number *R* of pieces in the piecewise demographic function needs to be specified. Setting *R* too low may lead to an oversimplified output that does not capture all the information on past population changes suggested by the genealogy, whereas setting *R* too high can lead to overfitting.

We therefore propose to use well established statistical methods to select the optimal value of *R*. First the model is fitted for multiple proposed values of *R*, and then for each output we compute the Akaike information criterion (AIC), which is equal to:

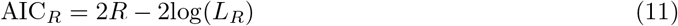

where *L*_*R*_ is the maximum value of the likelihood when using *R* pieces. The value of *R* giving the smallest value of AIC_*R*_ is selected. We also implemented the Bayesian information criterion (BIC), which is equal to:

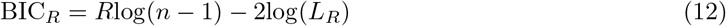

### Simulation of testing data

In order to test the accuracy of our methodology, we implemented a new simulator of coalescent genealogies given sampling dates and a past demographic function *N*_e_(*t*). When the demographic function is constant, the simulation of coalescent genealogies is equivalent to simulating from a homogeneous Poisson process, in which the waiting times from one event to the next are exponentially distributed. To extend this to the situation where the demographic function is non-constant requires to simulate from an equivalent non-homogeneous Poisson process. The approach we used to achieve this is to consider a homogeneous Poisson process with a population size *N*_m_ which is lower than any value of *N*_e_(*t*), i.e. ∀*t, N*_e_(*t*) ≥ *N*_m_. We simulate this process using exponential waiting times, but filter an event happening at time *t* according to the ratio *N*_m_*/N*_e_(*t*). Specifically, we draw *u* ∼ Unif(0, 1) and if *u < N*_m_*/N*_e_(*t*) the event is accepted and otherwise rejected. The resulting filtered Poisson process simulates from the non-homogeneous Poisson process as required (Ross 2014). The disadvantage of this approach over other methods of simulations is that there may be many rejections if *N*_e_(*t*) takes small values so that *N*_m_ needs to be small too. However, efficiency of simulation is not important for our purpose here, and this method has the advantage to avoid the computation of integrals on the *N*_e_(*t*) function which other methods would require.

### Implementation

We implemented the simulation and inference methods described in this paper into a new R package entitled *mlesky* which is available at https://github.com/emvolz-phylodynamics/mlesky. The optimisation of the demographic function makes use of the quasi-Newton Broyden-Fletcher-Goldfarb-Shanno (BFGS) method implemented in the optim command (Nash 2014). Confident intervals are computed based on an approximation of the curvature of the likelihood surface around its maximum. If multiple CPU cores are available, these resources are exploited within the procedure of selection of the smoothing parameter where the computation can be split between the different cross values in the cross-validation. Multicore processing is also applied in the procedure of selection of the grid resolution where computation can be split between different values of the resolution parameter *R*. All the code and data needed to reproduce our results on simulated and real datasets is available at https://github.com/mrc-ide/mlesky-experiments.

## RESULTS

### Application to simulated phylogeny with constant population size

A dated phylogeny was simulated with 200 tips sampled at regular intervals between 2000 and 2020, and a constant past population size function *N*_e_(*t*) = 20 (Figure S1). To illustrate the importance of the resolution *R* and precision *τ* parameters, we inferred the demographic function under the *skygrid* model (cf Equation 1) for a grid of values with *R* ∈ {5, 20, 50} and *τ* ∈ {1, 10, 20} (Figure 1). The results look quite different depending on the parameters used, and in particular when *R* is large and *τ* is small, fluctuation in the population size are incorrectly inferred. When applying the AIC procedure to this dataset, the correct value of *R* = 1 was inferred for which the parameter *τ* becomes irrelevant. In these conditions the effective population size was estimated to be 19.65 with confidence interval ranging from 17.10 to 22.57 which includes the correct value of 20 used in the simulation. We repeated the AIC procedure for 100 different phylogenies all which had been simulated under the same constant population size conditions described above. For 65 of these phylogenies the AIC procedure selected *R* = 1, with the third quartile falling on *R* = 3 and 94% of the simulations giving *R* ≤ 5. We also applied the BIC procedure for the same 100 phylogenies, and found that *R* = 1 was selected in all but one instance for which *R* = 2 was inferred. However, the BIC is well known to be overly conservative (Kuha 2004; Weakliem 1999) and so the rest of results make use of the AIC procedure.

**Figure 1:**
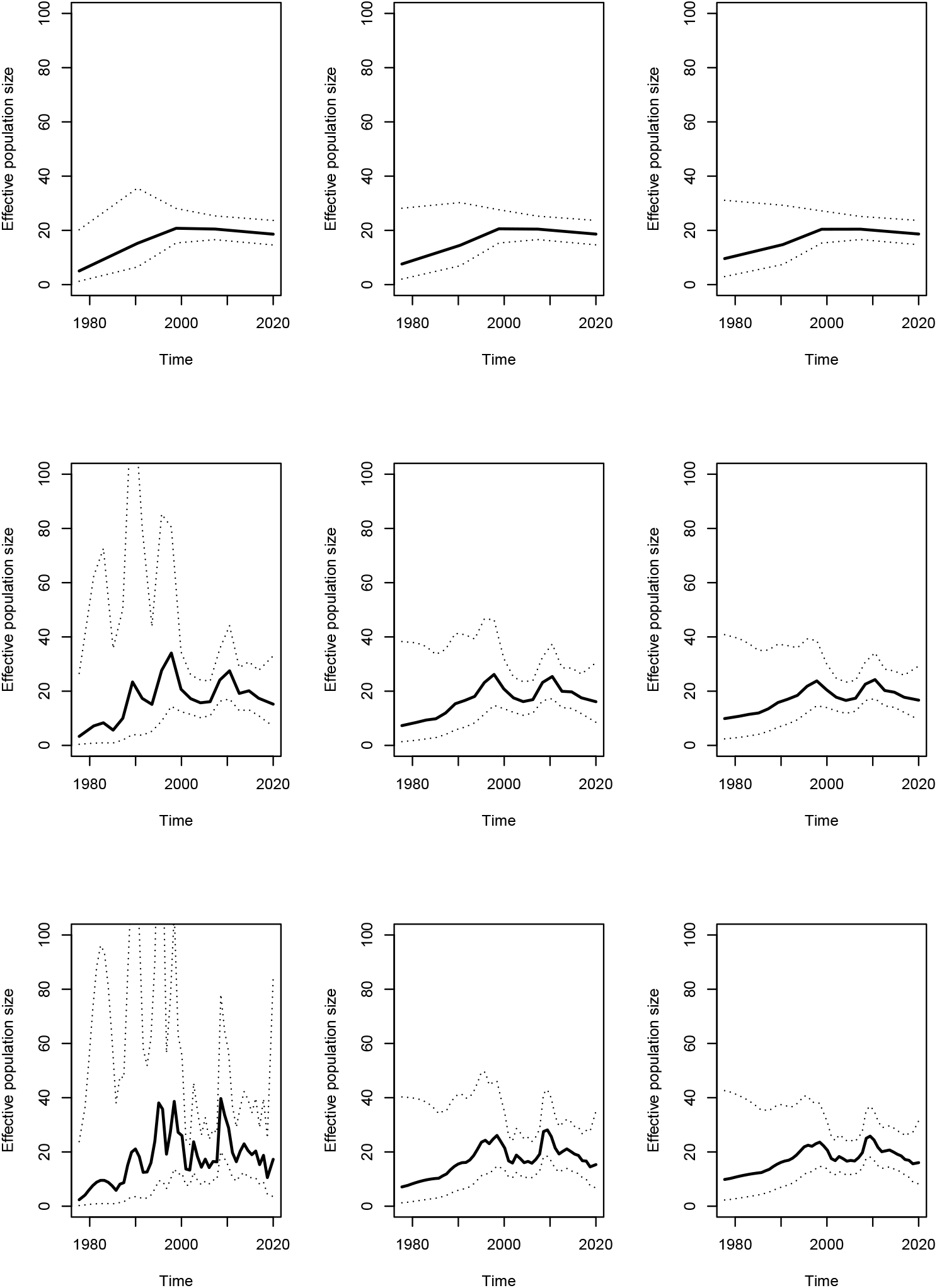
Result on simulated phylogeny shown in Figure S1 using the skyline model, from top to bottom *R* = 5, 20, 50 and from left to right *τ* = 1, 10, 20.

### Application to simulated phylogeny with varying population size

Next we simulated a dated phylogeny with the same number and dates of the tips as previously, but using a demographic function *N*_e_(*t*) that was sinusoidal with minimum 2 and maximum 22, with period 6.28 years. Figure S2 shows both the demographic function used and the resulting simulated phylogeny. We attempted to reconstruct the demographic function based on the phylogeny under the three models *skygrid, skygrowth* and *skysigma* described in Equations 1, 3 and 5, respectively. For each model the precision parameter *τ* was optimised using our new cross-validation procedure and the number of pieces was set to be *R* = 20 for ease of comparison. The results obtained in these conditions were very similar under the three models (Figure 2). This suggests that when the precision parameter is optimised using the cross-validation method, the choice between these three models becomes relatively unimportant. The same conclusions when reached when comparing the results of inference based on the three models to other simulated phylogenies. The choice of using one model rather than another is therefore mostly guided by the presence of covariate data and whether these are expected to correlate with the effective population size directly or some other function of it such as the population growth rates (Gill et al. 2016; Volz and Didelot 2018).

**Figure 2:**
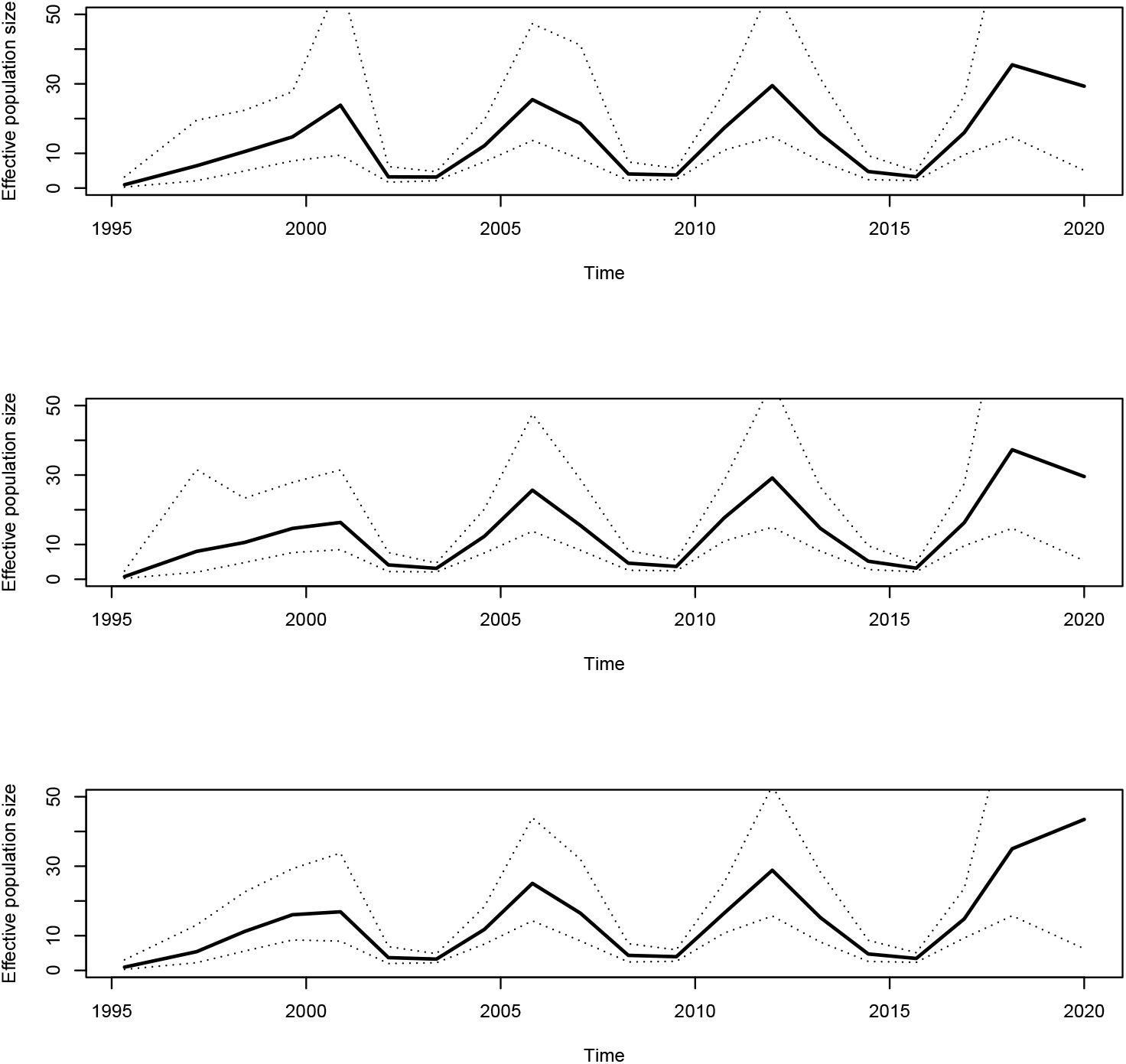
Result of applying the three different models (from top to bottom, *skygrid, skygrowth* and *skysigma*) to the phylogeny shown in Figure S2 which was simulated using a sinusoidal demographic function.

One situation in which all models are expected to perform poorly is when then there are sudden changes to the demographic function. To exemplify this, we simulated another dated phylogeny with the same and dates of the tips as before, but using a bottleneck function for *N*_e_(*t*) which was equal to 10 at all times except between 2005 and 2010 when it was equal to 1 (Figure 3A). The phylogeny simulated using this bottleneck function is shown in Figure 3B. We reconstructed the demographic function using the *skygrid* model. The lowest value of the AIC was obtained for *R* = 14, and the precision parameter was optimised using the cross-validation procedure to *τ* = 0.87. The inferred demographic function is shown in Figure 3C, where the bottleneck between 2005 and 2010 has been accurately detected.

**Figure 3:**
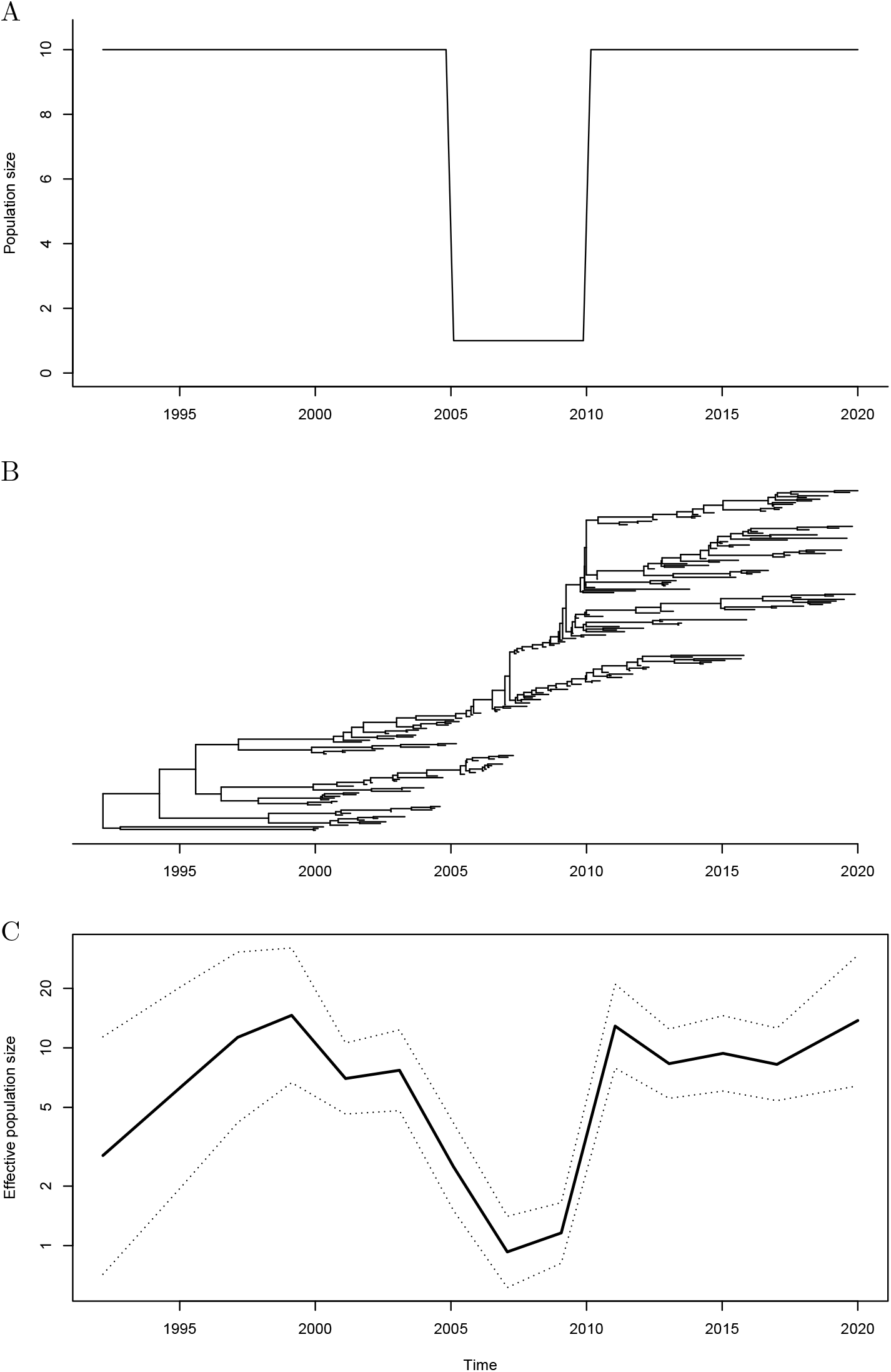
Demographic function (A), phylogeny (B) and inferred demographic function (C) for a simulated dataset under a bottleneck model.

### Application to simulated phylogeny with covariate data

Finally, we used simulations to test our procedure for the analysis of association between demography and covariate data. An example is shown in Figure S3 where the covariate data follows a simple quadratic function in order to create a boom and bust dynamic (Figure S3A). The growth rate of the population however does not follow exactly this function, and is subjected to monthly Gaussian noise with standard deviation 0.4 in this case (Figure S3B). From this growth rate we compute the effective population size function over time (Figure S3C) and simulate a phylogenetic tree as previously, with 200 tips sampled at regular intervals between 2000 and 2020 (Figure S3D). We then analysed this simulated phylogeny alongside the covariate data, and found in this case a strong association with coefficient *β* = 0.77. We repeated this procedure 100 times with increasing values of the noise standard deviation and the results are summarised in Figure S4. As expected, we found that as the noise increases, the coefficient of association *β* between growth rate and the covariate decreases, and eventually the association becomes non-significant with an estimated coefficient of association close to zero.

### Application to *Vibrio cholerae* dataset

We applied our methodology to a previously described collection of 260 genomes from the seventh pandemic of *Vibrio cholerae* (Didelot et al. 2015). A genealogy was estimated in this previous study using an early version of BactDating (Didelot et al. 2018), and it is reproduced in Figure 4A. We applied the AIC procedure to determine that the demographic function would be modelled using *R* = 16 pieces. The precision parameter was optimised to a value of *τ* = 1.84 using the cross-validation procedure. The whole analysis took less than 20 seconds on a standard laptop computer. The inferred demographic function is shown in Figure 4B. A first peak was detected in the 1960s, followed by a second peak in the 1970s and finally a third peak in the 1990s. This demographic function follows closely on the previously described three “waves” of cholera spreading globally from the Bay of Bengal (Mutreja et al. 2011; Didelot et al. 2015; Weill et al. 2017). However, these three waves had previously been described based on phylogeographic reconstructions of the spread of the pandemic around the world. The fact that we found a similar wave pattern in our analysis which did not include any information about the geographical origin of the genomes provides further support for the validity of this phylodynamic reconstruction.

**Figure 4:**
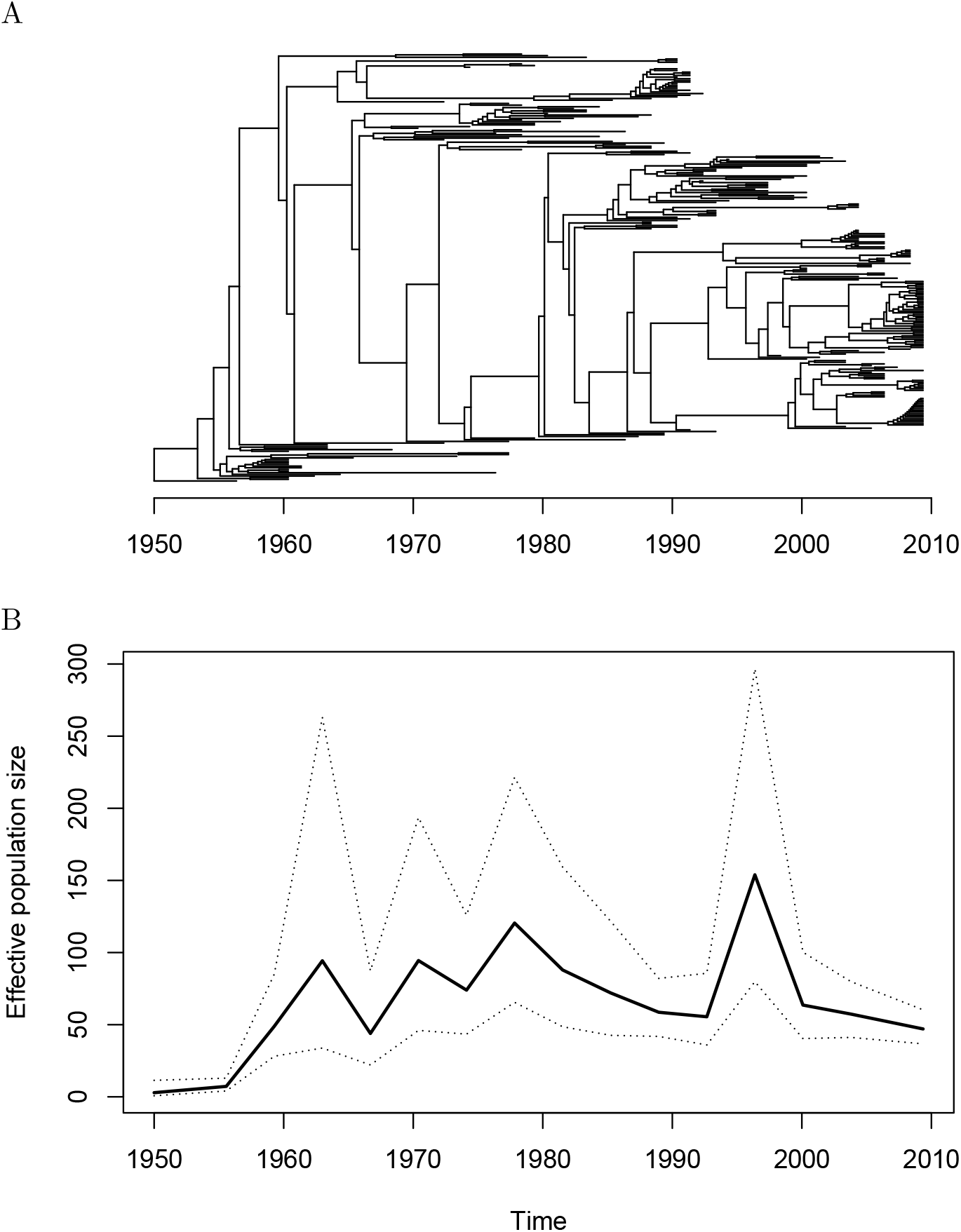
Analysis of the seventh pandemic of *Vibrio cholerae*. (A). Dated phylogeny used as the starting point of past population size inference. (B). Demographic function reconstructed based on the phylogeny above.

### Estimating the impact of non-pharmaceutical interventions for COVID-19 in England

We applied our methodology to the SARS-CoV-2 epidemic in England using data from the first epidemic wave spanning the spring of 2020. By incorporating data on timing of public health measures such as lockdowns, we estimated the association on non-pharmaceutical interventions (NPIs) with viral transmission. The COVID-19 Genomics UK Consortium (COG-UK) was established on 23rd March 2020 and has coordinated a large-scale sequencing and bioinformatics effort to assist with COVID-19 surveillance and response (COG-UK Consortium 2020). The proportion of cases with a virus genome has varied over time and increased rapidly in April 2020 following the establishment of large-scale national sequencing laboratories. In order to facilitate molecular clock dating, we carried out a stratified random sample of genomes between 1st January and 30th April 2020 ensuring good representation of sequences across a wide range of calendar time. Sequences were ordered by sample date, binned by day, and randomly selected from each bin. Duplicate sequences were removed. Repeating this process ten times resulted in ten distinct sequence sub-samples with a mean of 4,217 sequences each.

As part of the COG-UK bioinformatics pipeline, a maximum likelihood tree is estimated at regular intervals on the MRC-CLIMB infrastructure (Nicholls et al. 2021). We pruned these trees to retain samples in each of our sequence sub-samples. Each of these sub-trees was then converted into time-scaled phylogenies using treedater v0.5.1 (Volz and Frost 2017) by randomly resolving polytomies in the tree and sampling a molecular clock rate of evolution from a normal distribution with mean 5.91 × 10^−4^ substitutions per site per year and standard deviation 1.92 × 10^−5^, based on previous analysis of SARS-CoV-2 in the UK (Volz et al. 2021). In all, 100 time trees were estimated representing uncertainty in phylogenetic dating and sampling variation. The *skysigma* model was fitted to the trees by maximizing the combined (average) likelihood. Maximum likelihood estimates were also computed for each tree and 95% quantiles were used to quantify the uncertainty in the parameter estimates. The results are shown in Figure 5A for the estimation of the effective population size function and in Figure 5B for the estimation of the basic reproduction number over time. The latter is calculated as *R*(*t*) = *ρ*(*t*)Ψ + 1 where *ρ*(*t*) is the growth rate of the effective population size *N*_e_(*t*) estimated through time and Ψ is the mean of the serial interval (Wallinga and Lipsitch 2007; Volz and Didelot 2018). The value Ψ = 6.5 days was used based on previous studies of infector-infectees pairs (Chan et al. 2020; Bi et al. 2020; Wu et al. 2020).

**Figure 5:**
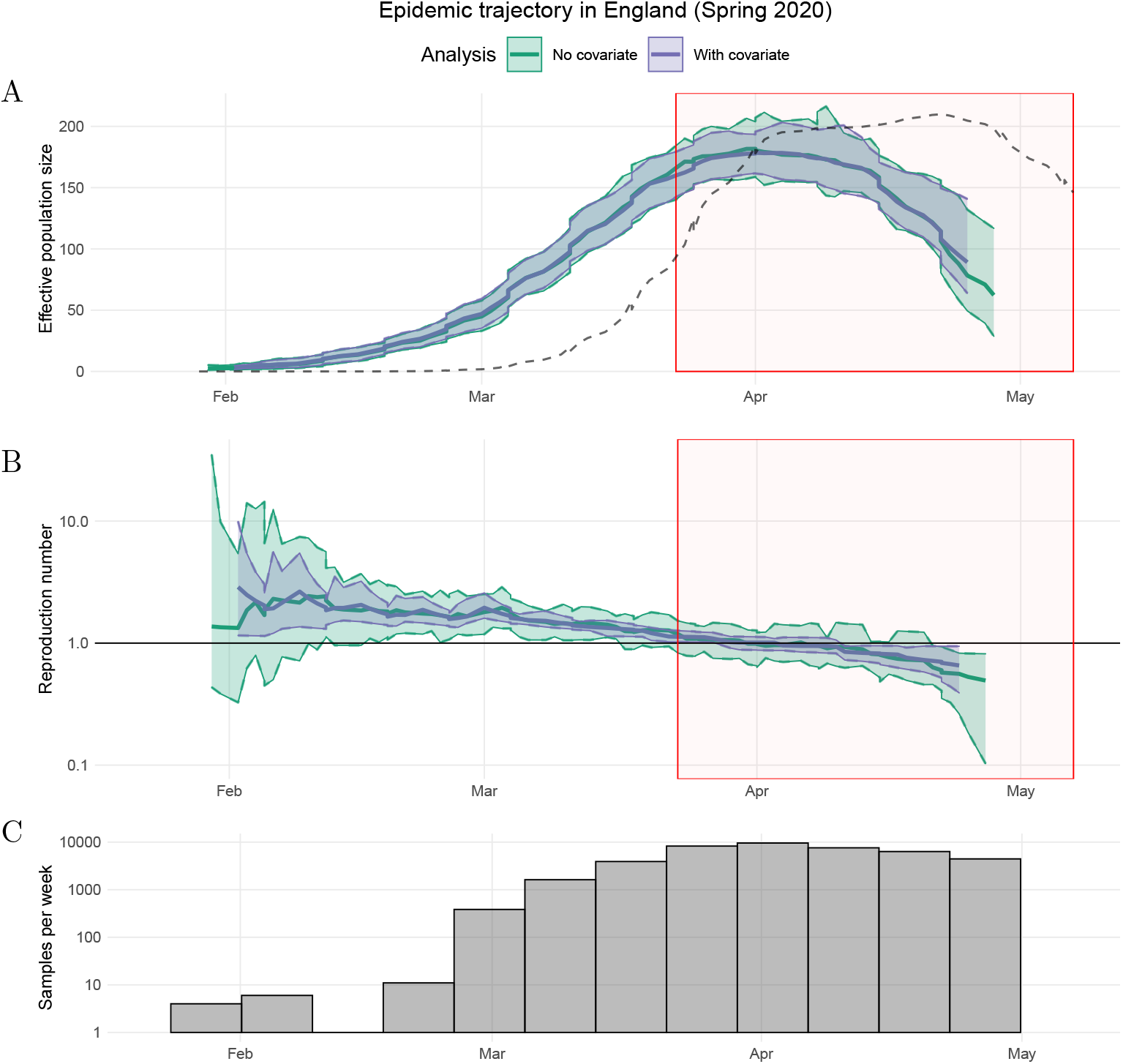
The epidemiological trajectory of SARS-CoV-2 in England during spring 2020. Thick solid lines and shaded areas represent the median and 95% quantiles of *N*_e_(*t*) with (purple) and without (green) the OxCGRT health containment index as a covariate of *N*_e_(*t*) growth rates. The model is fitted with no back-shift in the covariate. Red shaded area represents period of first national lockdown in England. Black dotted line represents daily confirmed cases (smoothed and rescaled). (A) Effective population size *N*_e_(*t*) through time. (B) Reproduction number *R*(*t*) through time. (C) Frequencies of sample dates for tips in each sample week in the SARS-CoV-2 phylogenies.

The estimated peak of the epidemic occurred on 1st April 2020, eight days after the imposition of the first national lockdown, illustrated by the red boxes in Figure 5. The rise and fall in *N*_e_(*t*) precedes a similar dynamic in the number of confirmed cases by several weeks (Figure 5A), which is as expected since the case ascertainment rate was initially very low and improved dramatically in April. On the other hand, the estimated *N*_e_(*t*) is approximately consistent with the number of genomic sequences available over time (Figure 5C). The estimated *R*(*t*) decreased gradually in the three weeks preceding the start of the national lockdown (Figure 5B). This may be due to changing behaviour prior to the national lockdown and a changing proportion of cases due to travel-related importation. Travel-linked cases declined while internal transmission increased throughout March and April (du Plessis et al. 2021).

To test for association between growth rates and NPIs, we also fitted the model to both the genealogical data and the OxCGRT health containment index (Hale et al. 2020), a time series representing the intensity of the public health response. A higher value of this index indicates more stringent NPIs. The model was fitted under the assumption that the differential of the logarithm of *N*_e_(*t*) follows differential of the OxCGRT index, which approximately corresponds to the hypothesis that the basic reproduction number *R*(*t*) follows the daily change in the index (Volz and Didelot 2018). The median estimated epidemic trajectories are very similar when including this covariate (Figure 5A), and we observe improved precision in the estimate of the reproduction number (Figure 5B).

Figure 6 shows the coefficient *β* which represents the estimated strength of association between the reproduction number and the daily change in the OxCGRT index. A negative value of *β* indicates a negative association between changes in NPIs and the reproduction number. We investigated how the estimate of *β* depends on a lag (back-shift) between the value of the index and the demographic function. The largest effects (most negative values) were found when shifting the index back 8 days, which means relating the values of the index to the reproduction number 8 days later. In contrast, when changes in NPIs are compared to growth rates that precede them by several days (negative delay), the coefficient *β* is not significant.

**Figure 6:**
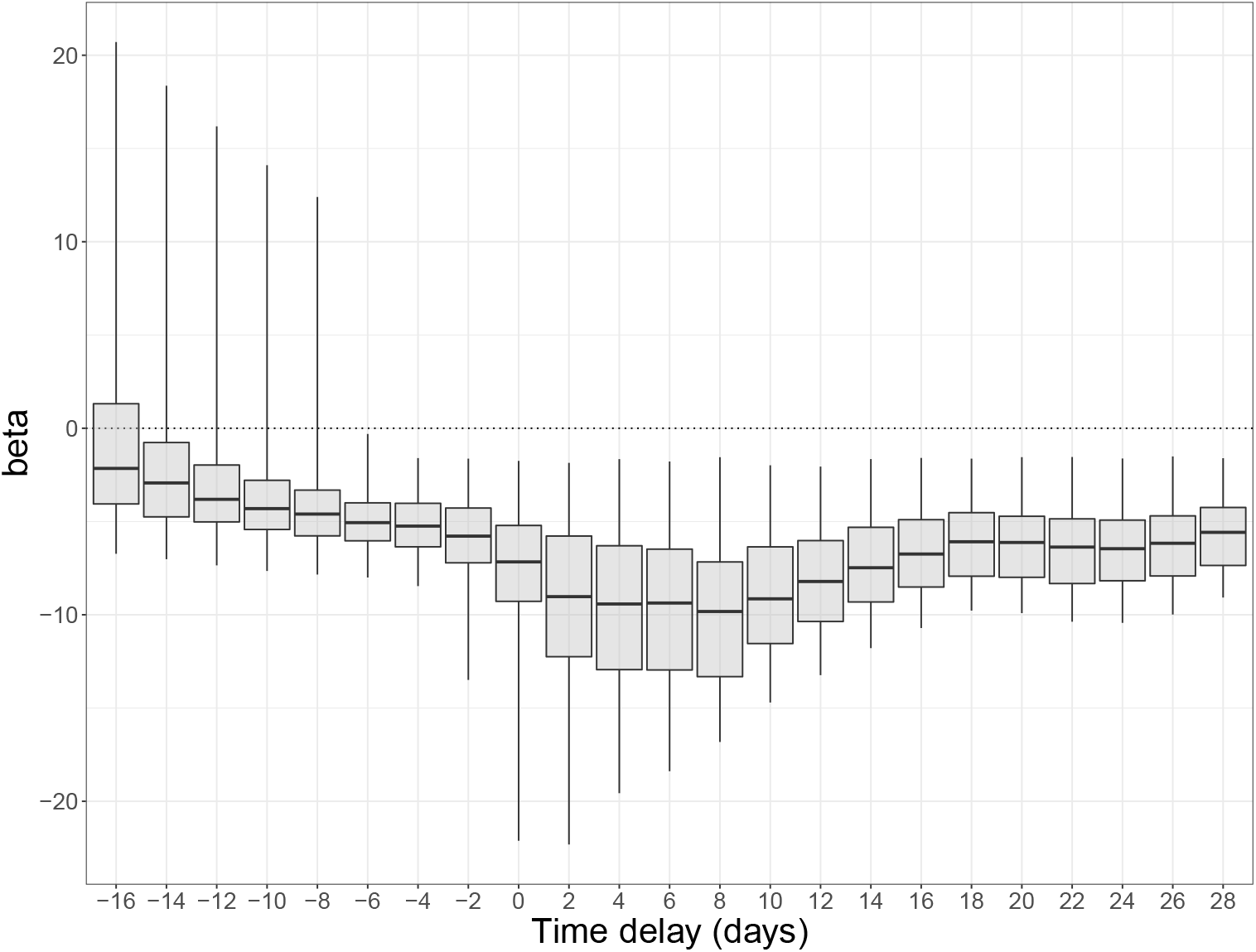
Distribution of association coefficients when testing for univariate association between daily changes in the OxCGRT health containment index and daily changes in the reproduction number of SARS-CoV-2 in England. Boxes represent the median and interquartile range; whiskers show 95% quantiles. A positive delay of 10 represents testing an association between the OxCGRT index at time *t* and the reproduction number at time *t* + 10 days.

## DISCUSSION

Non-parametric phylodynamic inference of population size dynamics is usually carried out in a Bayesian framework (Drummond et al. 2005; Minin et al. 2008; Gill et al. 2013). Here we presented methods for performing such inference in a frequentist setting with a particular view towards model selection and avoiding over-fitting. Optimal smoothing can be obtained in a natural way using standard cross-validation methods, and the optimal resolution of the discretised demographic function is achieved using the well-established AIC criterion. This approach can be advantageous when prior distributions are difficult to design or results are sensitive to arbitrarily chosen priors. Methods based on likelihood maximization are also fast and scalable to datasets much larger than is conventionally studied with Bayesian methods, and the selection of smoothing parameters does not require arbitrarily chosen hyperparameters. Conventional AIC metrics also alleviate the difficulty of model selection. In most of our simulations, we find relatively little difference in our estimates when parameterizing the model in terms of log(*N*_e_(*t*)) (Equation 1), the growth rate of *N*_e_(*t*) (Equation 3) or the second order variation of log(*N*_e_(*t*)) (Equation 5), as long as the precision parameter *τ* for each model is optimized as we proposed.

Our methodology assumed that a dated phylogeny has been previously reconstructed from the genetic data. It is therefore well suited for the post-processing analysis of the outputs from *treedater* (Volz and Frost 2017) or *TreeTime* (Sagulenko et al. 2018). A key assumption of our method, as with its Bayesian counterparts, is that all samples in the phylogeny come from a single population ruled by a unique demographic function. To ensure that this is indeed the case, complementary methods are emerging that can test for the presence or asymmetry or hidden population structure in dated phylogenies (Dearlove and Frost 2015; Volz et al. 2020). Conversely, if multiple phylogenies follow the same demographic dynamic, they can be analysed jointly to provide a more precise reconstruction of the demographic function and epidemiological parameters (Xu et al. 2019), and our software implementation is able to perform such a joint analysis when appropriate.

Past variations in the effective population size of a pathogen population can reveal key insights into past epidemiological dynamics and help make predictions about the future. It is important to note that the effective population size is not generally equal to or even proportional to the number of infections over time (Volz et al. 2009; Dearlove and Wilson 2013). On the other hand, the growth rate of the effective population size can be used to estimate the basic reproduction number over time *R*(*t*) (Wallinga and Lipsitch 2007; Volz et al. 2013; Volz and Didelot 2018) as we used in our application to COVID-19 in England. Having good estimates of this quantity is especially important for assessing the effect of infectious disease control measures (Fraser 2007), and phylodynamic approaches provide a useful complementary approach to more traditional methods of estimation based on case report data (Cori et al. 2013).

## Acknowledgements

XD acknowledges funding from the National Institute for Health Research (NIHR) Health Protection Research Unit in Genomics and Enabling Data. LG acknowledges funding from the MRC Doctoral Training Partnership. EMV was supported by NIH R01-AI135970. EMV acknowledges the MRC Centre for Global Infectious Disease Analysis (MR/R015600/1). All authors thank the COVID-19 UK Consortium for providing data and bioinformatics resources. We thank all partners and contributors to the COG-UK consortium who are listed at https://www.cogconsortium.uk/about/. COG-UK is supported by funding from the Medical Research Council (MRC) part of UK Research & Innovation (UKRI), the National Institute of Health Research (NIHR) and Genome Research Limited, operating as the Wellcome Sanger Institute.

---

**Figure S1:**
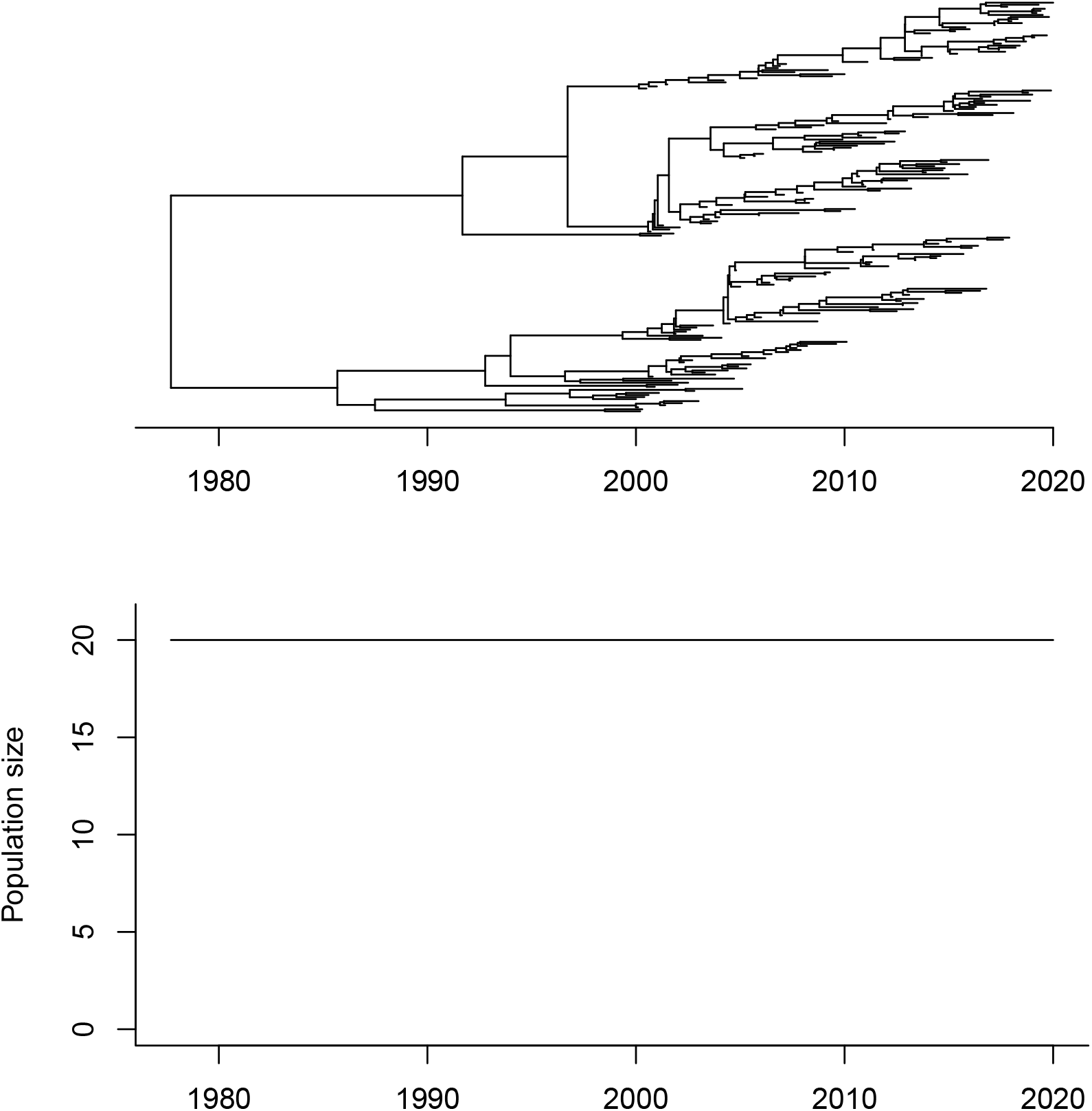
Simulated phylogeny using a constant demographic function.

**Figure S2:**
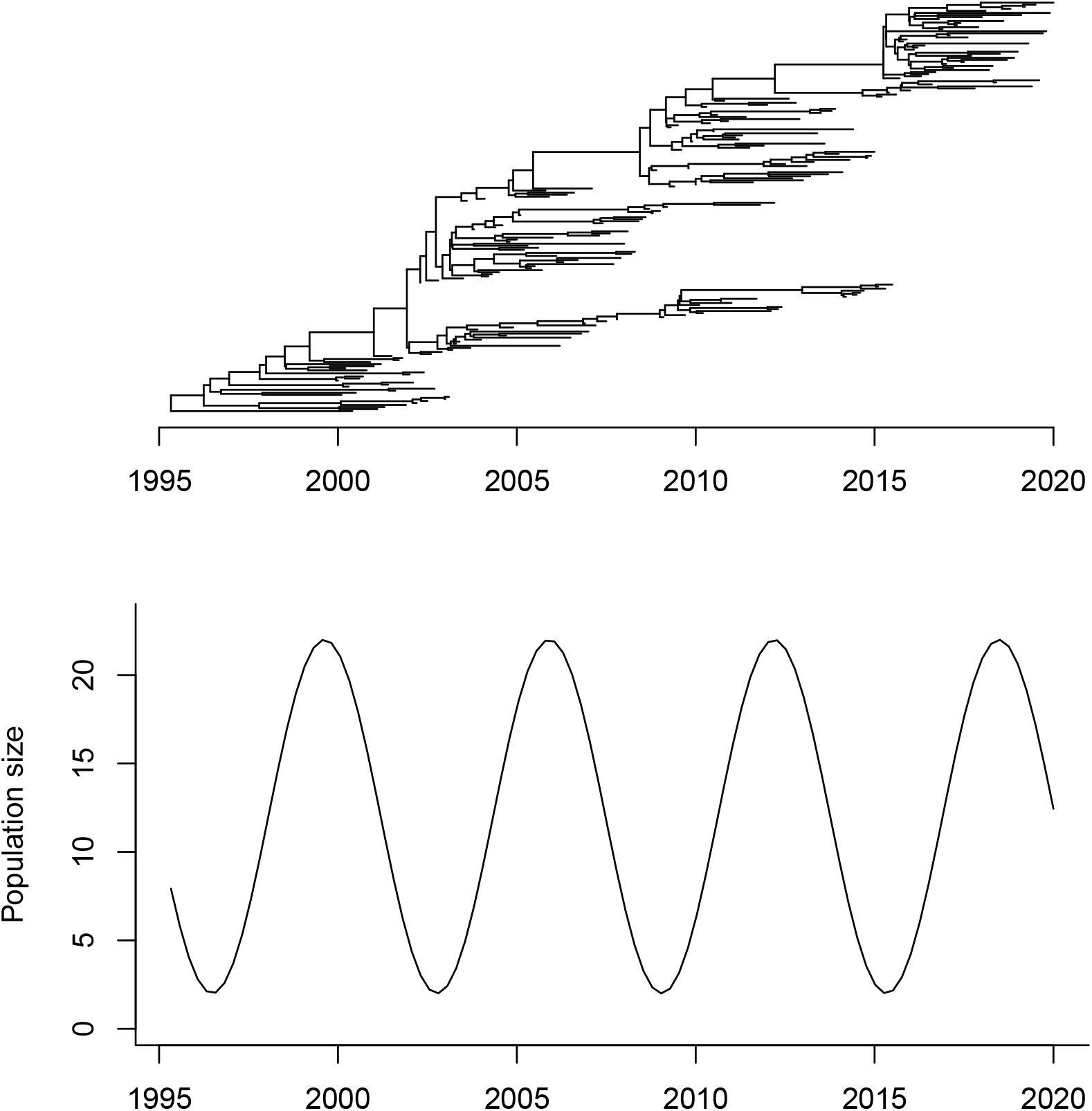
Simulated phylogeny using a sinusoidal demographic function.

**Figure S3:**
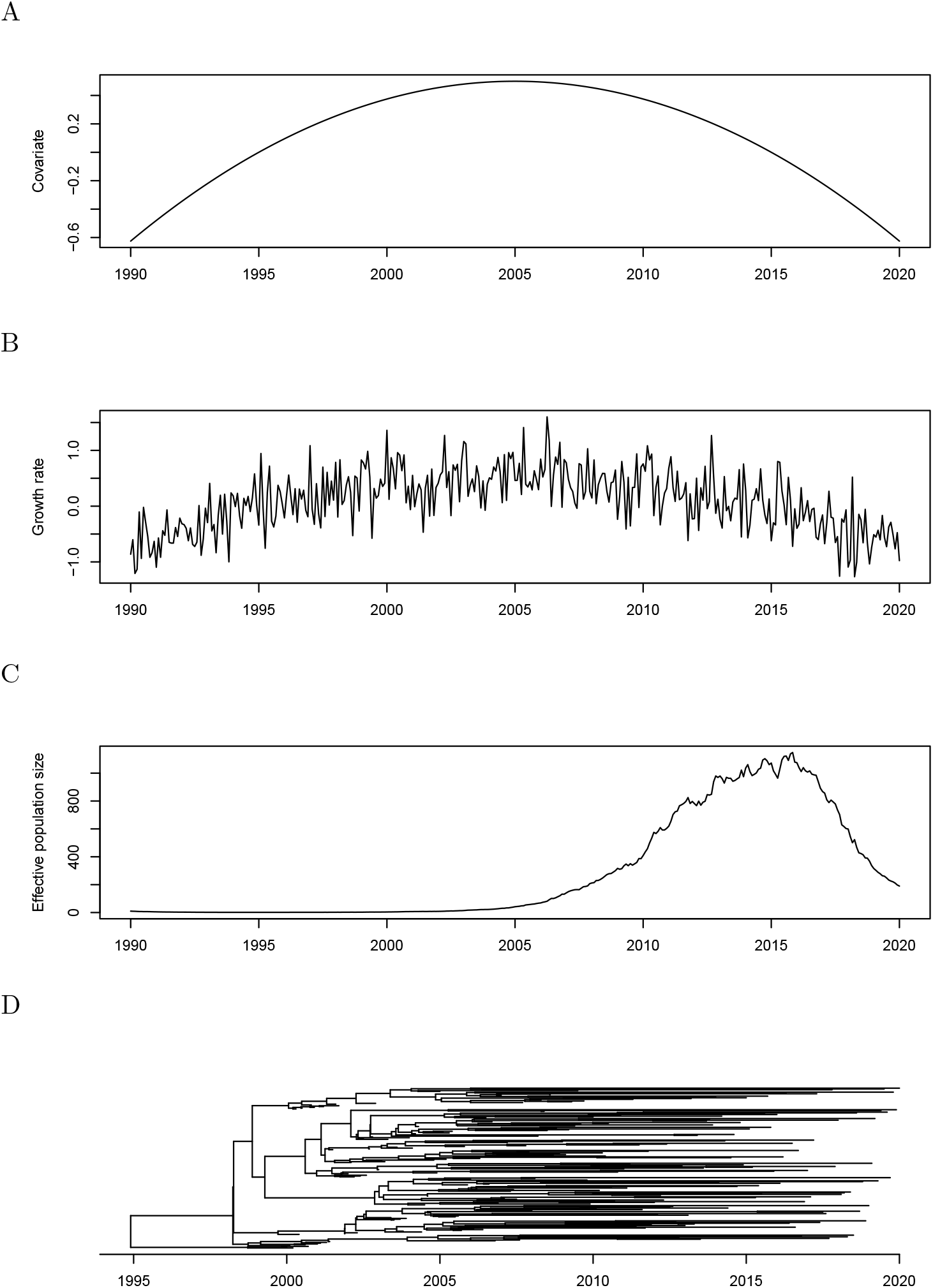
Example of simulation with covariate data driving the growth rate. (A) Covariate data following a quadratic function. (B) Growth rate equal to the covariate data plus some Gaussian noise. (C) Effective population size. (D) Dated phylogeny.

**Figure S4:**
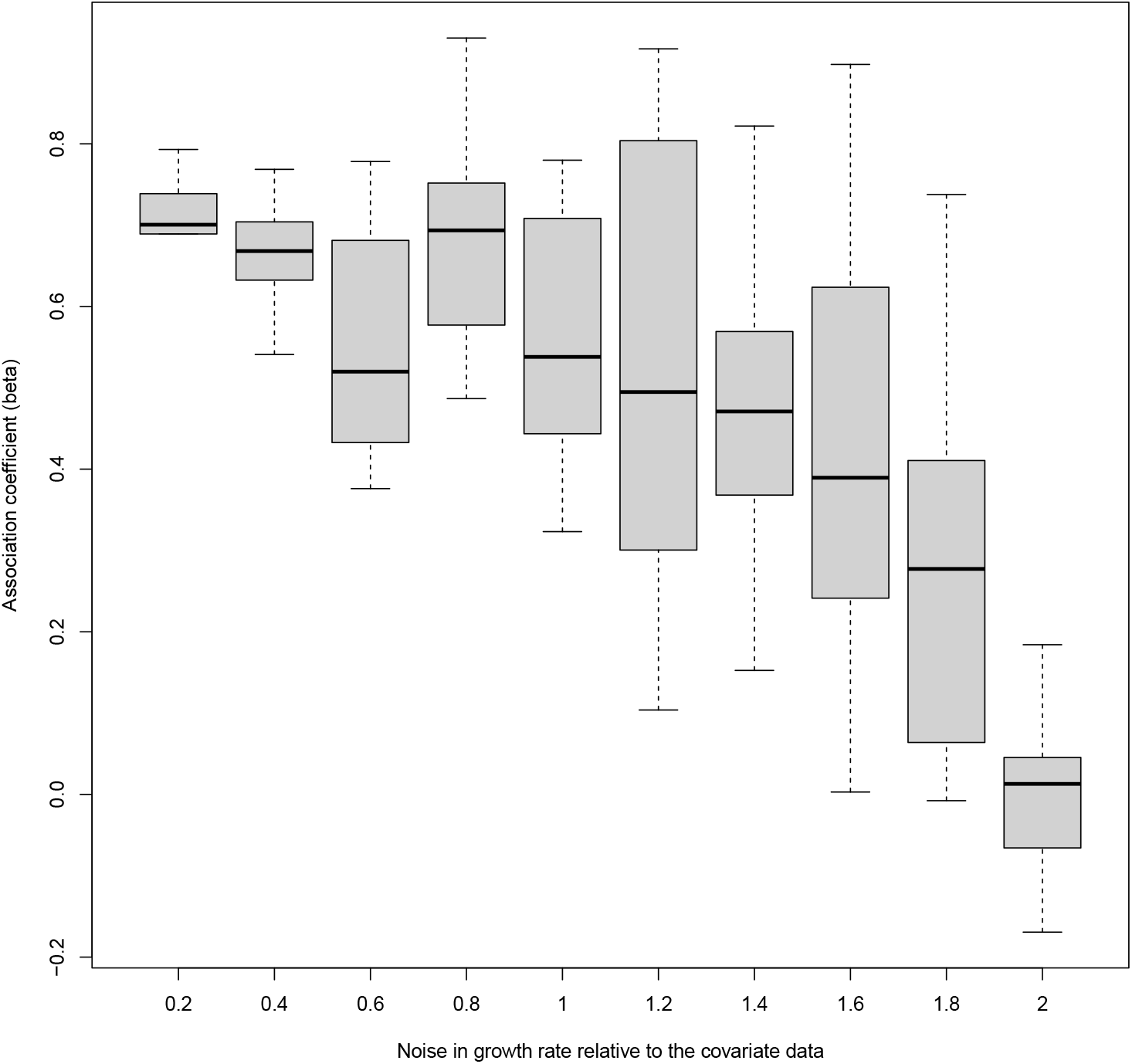
Results of the covariate analysis. For each value of the Gaussian noise (x-axis) ten simulations were performed and the inferred values of the association coefficient *β* are shown (y-axis) as boxplots.

## Notes

### Competing Interest Statement

The authors have declared no competing interest.

